# Hidden rAAV Breakpoints Detected Using Single-Molecule, Modified Base Sequencing

**DOI:** 10.1101/2023.11.06.565870

**Authors:** Terrence Hanscom, Luis M M Soares, Alice Zheng, Nathanael Bourgeois, Katherine Gall, Thia St Martin, Jason Wright, Donald E Selby

## Abstract

The AAV genome is a single stranded DNA molecule packaged in an icosahedral protein capsid. Vector genomes of plus and minus polarities are packaged and complementary genomic strands hybridize when lysed *in vitro*. Standard sequencing library methods cause loss of information from individual genomes when mismatches and gaps are repaired. To retain original molecular information, modified bases are used during the repair step which allows pre-existing DNA to be distinguished from DNA added during library preparation. Modified bases introduced during repair are identified using the Sequel II system and used to detect HIDdEN DNA breakpoints (HIDEN-Seq). The most frequent breakpoints in an AAV vector subject to high strand breakage during packaging were linked to adjacent secondary structure, prompting changes in nearby sequences to reduce breakage. This use of modified bases for localizing DNA breaks enables better vector design, resulting in higher quality gene therapy vectors. The same approach can be used in other systems where knowledge of pre-existing sequence and structure is important.

## Introduction

Recombinant Adeno-Associated Virus (rAAV) is an important gene therapy vector due to its low immunogenicity and long track record for safe use in humans (1, 2). As with all gene therapy systems, the quality of the therapeutic material delivered is critical for both safety and efficacy (3), but AAV’s genomic properties make its detailed characterization challenging. AAV packages its genome into pre-assembled capsids using the viral protein Rep which is responsible for cleaving packaged DNA at the correct position (Fig 1) (4). This process can result in non-functional, empty capsids and partially filled capsids or the intended completely full capsids.

**Figure 1.**
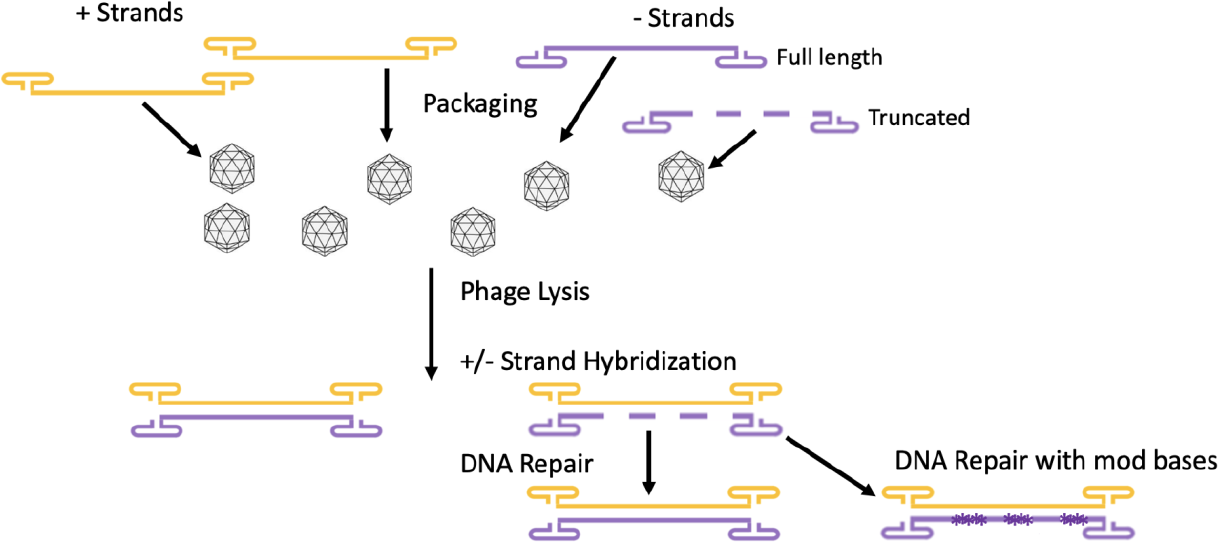
Cartoon Simplification of Packaging and hybridization of AAV ssDNA

The presence of empty or partially full capsids lowers viral titer, negatively affecting rAAV quality, increasing manufacturing costs, and increasing the likelihood of adverse effects during treatment. A variety of methods have been used to measure the relative amounts of the different products (5), but they typically provide little detail about the exact nature of DNA in the capsids and no insight into improving the yield of full capsids.

Typical results obtained when sequencing capsid DNA can be misleading for multiple reasons. When short-read sequencing is used, the entire genome is not provided within single reads, so overlapping reads are of unknown continuity. Long-read sequencing can overcome this problem; but, using standard methods, it cannot readily distinguish the sequence differences from the opposite strands of ssDNA that come from different capsids. AAV is packaged as single-stranded DNA and when the capsids are lysed *in vitro* and DNA is released into solution, DNA of opposite strandedness can hybridize into heteroduplexes even if the DNA is not full-length. Using standard library protocols, DNA is “repaired” and then acted upon by ligases or transposases to add sequencing adaptors. While repair of incidental DNA damage is useful for many applications, some repairs eliminate pre-existing, real DNA features including potentially important sequence and length variations present in the original AAV or other DNA. When AAV containing both partial and full-length genomes is processed, the original breakpoints and gaps are “repaired” so both the number and sequence of the original truncated molecules are hidden and the DNA falsely appears to be full-length.

To eliminate the problem of DNA repairs concealing real sequence features or propagating artifacts (6), we developed a method that allows us to distinguish pre-existing DNA from DNA added during the library preparation. Instead of using four standard nucleotides, two (or one) methylated nucleotides are replaced into the library preparation so that any new DNA added during repair includes stretches of modified bases where there had previously been sequence gaps while the pre-existing ssDNA retains its natural, unmodified state. Using the PacBio Sequel II system, modified bases can be distinguished from unmodified bases (7-9), enabling breakpoint identification at high resolution.

Incorporation of modified nucleotides during the DNA repair process allows us to study rAAV samples known to contain truncated genomes caused by breaks that occur during packaging, a phenomenon that is found in all AAV serotypes. Being able to separate DNA inserted during repairs from pre-existing DNA packaged in capsids allows us to map the sites that were fracturing and more accurately measure the packaged lengths of rAAV ssDNA molecules. These rAAV breakpoint data have allowed us to better understand the causes for DNA breakage, to design better therapeutic vectors, and to generate more precise quality control data.

## Results

While multiple approaches have been used to analyze partially filled AAV capsids and the resultant truncated DNA, previous methods do not provide insight into the detailed nature of the truncated molecules. Alkaline agarose gels can be used to visualize the length of DNAs from capsids (10) though their low resolution and lack of sequence or strand information makes them less useful for establishing exact breakpoints within the DNA. Individual samples can have widely divergent levels of full-length versus truncated genomes (Fig 2A). For example, sample 1 has less full-length DNA as a fraction of the total than sample 2 that is primarily full length.

**Figure 2.**
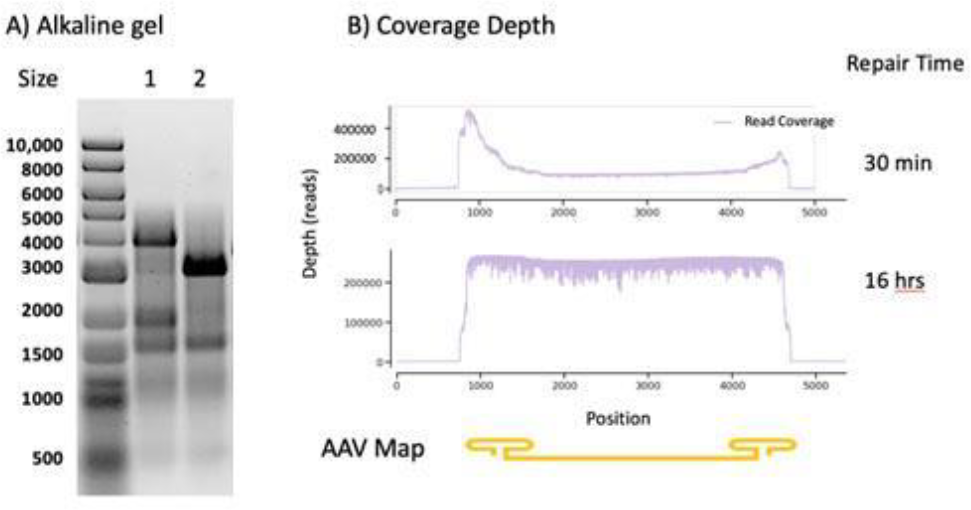
Genomic DNA size and sequence coverage

However, the coverage depth plot produced by standard sequencing for sample 1 (Fig 2B) does not reflect the low level of full-length DNA. This discrepancy between sequencing and gel sizing can be explained by the repair processes used during standard NGS library preparation protocols. Gaps are filled and ligated to adjacent DNA, obscuring the original state of the DNA. A confounding issue was an end bias that was also observed with the standard DNA damage repair time of 30 minutes. This can be seen on the (upper graph Fig 2B) the standard DNA damage repair time of 30 minutes resulted in an end bias in the read coverage. This was alleviated by increasing the DNA damage repair time to 16 hours (lower graph Fig 2B).

To better assess the length and sequence of AAV DNA, it is necessary to use long-read sequencing to capture the full-length sequence as a continuous read and be aware of the impact of DNA repair on sequence output. The ITRs at both ends of the molecule are particularly problematic because of their high degree of secondary structure and GC-richness (11). We have found that the most accurate sequencing of ITRs and the remainder of the AAV genome is achieved using the Pacific Biosciences (PacBio) Sequel II in the circular consensus sequencing (CCS) mode (12). This method provides real-time, long-read, single molecule sequence data with multiple reads of each DNA so that a robust consensus sequence can be generated (13). The data can also be segregated by strand, allowing independent assessment of each strand as individual molecules with their own sets of sequence information.

For base-calling, the identity of each base is established by detecting which of four colored fluorophores is bound to a tethered polymerase prior to incorporation into the extending DNA. In addition to the color data that marks the identity of the incorporated base, there is also kinetic information about the Pulse Width (PW, time in which the incoming nucleotide is bound to the polymerase prior to incorporation) and the Inter-Pulse Distance (IPD, time between the incorporation of one nucleotide and the binding of the next one) (Fig 3). Standard nucleotides have a natural variance in these numbers, but modified bases can have substantially different values that readily allow their distinction from standard bases. We chose two commercially available modified nucleotides, 4-methyl-dCTP and 6-methyl-dATP because of their distinct impact on PW and IPD that allows their ready detection during sequencing (7). To maximize sensitivity of detection, CCS mode was used where the same molecule is sequenced multiple times in order to improve accuracy of base-calling, PW, and IPD. As shown in Fig 3, both the PW and the IPD near the modified base can be substantially longer. There is a more dramatic difference in IPD versus PW for the modified bases used here so we have employed that measure in these experiments. To maximize the sensitivity of the measurements, we require that at least 3 cycles are available for each strand analyzed. With these thresholds, we detect HIDdEN DNA breakpoints and we refer to this method as HIDEN-Seq.

**Figure 3.**
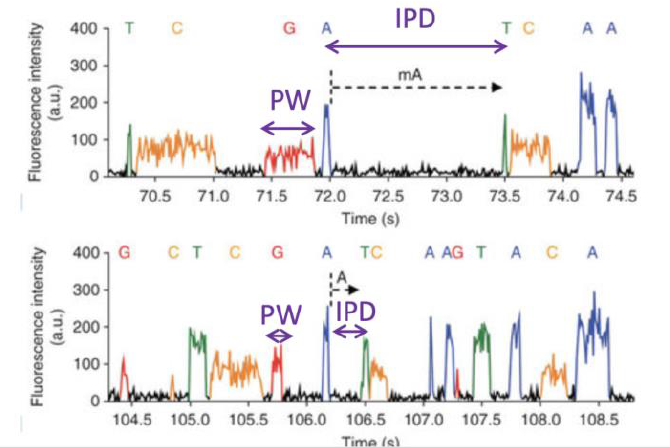
Variation of IPD and PW

As shown in Fig 4, the number of modified base patches induced by hidden breaks correlated with the fraction of nicked DNA. This indicates that the fraction of truncated or broken DNAs can be estimated using the number of patches of modified DNA.

**Figure 4.**
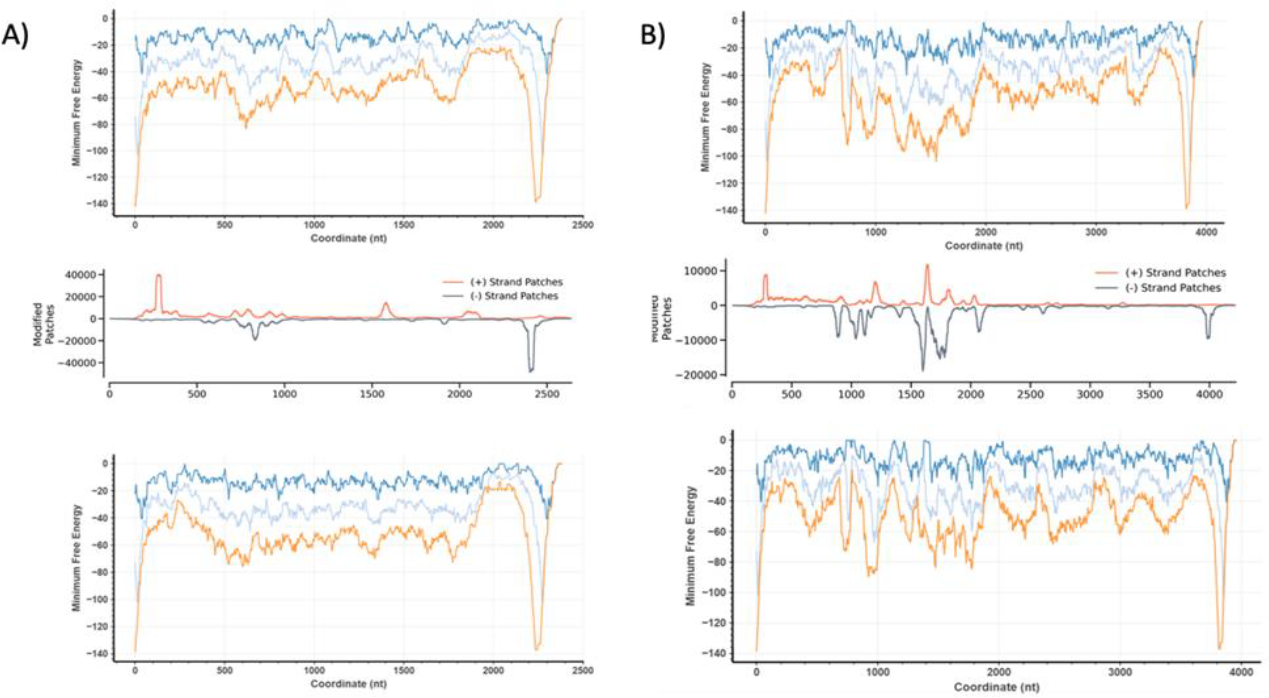
Modified Base Patches and Secondary Structure as a function of position

We chose an AAV sample known to have an issue with truncated genomes based on standard quality measurements (14). When subjected to the normal library preparation methods, the sequence data did not recapitulate the high degree of truncation observed using alkaline agarose gel electrophoresis, suggesting that the strand breaks were being obscured by sample processing (Fig 2). To address this, we replaced the standard nucleotides found in most DNA repair kits with two standard (dTTP and dGTP) and two modified nucleotides (4-methyl-dCTP and 6-methyl-dATP). A variety of polymerases, concentrations, and ligase ratios were tested to maximize the ability to fill gaps within the DNA with modified bases while minimizing elimination of pre-existing DNA via exonuclease or other activities. A high ratio of ligase to polymerase is used to seal gaps once they have been filled by the polymerase.

Each double-stranded AAV genome that is sequenced can arise in at least two different ways. The ITR at the 3’ end of the genome provides both a double-stranded region and a free 3’ OH so that site can potentially self-prime the AAV vector to generate a partial or complete second strand copy using the original ssDNA as a template. The use of modified nucleotides allows this to be easily distinguished from the contrasting situation in which two single-stranded AAVs of opposite strand polarity anneal. This hybrid molecule can potentially provide unique sequences on each strand arising from differences in the original molecules. As long as those sequence differences are not removed by repair protocols, the sequence of each can be determined independently. No amplification of DNA is carried out in our studies, so the single-molecule read count, broken down by strand, is a reflection of the frequency of each molecular species. With tens of thousands of reads for each sample, the frequency of truncations is determined by the number of patches of modified bases (Fig 4). Modified bases are detected by comparing the observed IPD at each position relative to what it would be without modification, using the metric IPD ratio, which is the ratio of the mean IPD at a site in the native sample to the mean IPD at the same site in the amplified control. Based on the distribution of IPD ratios for standard and modified bases established by Zatopek et al., we resolved the threshold for identifying a modified base to having an IPD ratio of above 3.0 (9). We define a patch as a region or base with a high concentration of modified bases. Each patch of bases that is defined as modified arises from the action of repair enzymes that insert modified bases in that region. Those repairs could be due to a gap in the sequence arising from strand breaks or from a free 3’ OH as would be found at the naturally occurring cleavage site at the AAV 3’ end. The location of the patch can be used to infer how it might have arisen.

The frequency of break points varies significantly by vector and is often highly localized to particular locations. Inspection of those locations shows that many contain a high level of sequence-predicted secondary structure. The predicted secondary structure strength for the tested vectors is shown in Fig 4 based on the Vienna RNA secondary structure algorithm (15). All the most frequent truncations occur close to the locations of the strongest secondary structures. The threshold for high level truncations occurs around -80 kcal/mole for a 150 nt segment. Hairpins stronger than that cleave more often than the ITRs while weaker hairpins cleave less frequently. When sequence modifications are made to these regions, the truncation frequency drops (Fig 4).

## Discussion

AAV presents unique challenges in its analysis due to its packaging as individual single strands with both orientations present in different capsids. Upon capsid lysis, inter-DNA hybridization is virtually unavoidable, generating a mixed DNA containing sequence and length information from two or more viral particles. This can lead to mischaracterization of what is actually present in the original viral mix because the gaps are filled in and can be ligated to other DNAs during the standard repair process. Similar loss of sequence information has also been observed by others using artificial systems (6).

With virtually all library preparations for DNA sequencing, there are DNA repair steps, DNA replication/amplification reactions, and/or ligations/molecular manipulations. These steps can introduce changes into the starting DNA that impact the sequence output and can change the observed sequence relative to the original input DNA though there is a published method (16) in which the DNA is sequenced directly without any modifications or additions. For the most part, these DNA modifications improve sequence quality and do not negatively impact the final analysis, but they can lead to a variety of different artifacts or misinterpretations. For example, artificial fusions can be generated by ligation of unrelated DNA fragments in a complex cellular mix and be mistaken for DNA integrations or structural rearrangements. Use of proper controls allows these to be resolved (17, 18), but it is not uncommon for these artifacts to be misinterpreted. Similarly, sequencing AAV ssDNA can lead to DNA-repair induced artifacts.

When there is a desire to know exactly what was in the starting DNA mixture prior to repair and sequencing, it is important to understand how the original DNA was manipulated during the library preparation and sequencing process. This is the certainly the case for DNA used in gene therapy applications where quality control is a key issue (19). There are other sequencing applications where this may be critical. However, there are other applications, such as genome assembly, that are minimally impacted by manipulations carried out during DNA processing or those steps have a positive effect on sequence quality outcome.

Previously, detection and localization of modified bases has been used for mapping nucleosomes, protein binding sites, and replication forks (8, 20-23). We have extended the use of modified bases for inclusion during the repair stage of library preparation. This allows distinction of the pre-existing, standard bases originally present in the viral capsid DNA relative to the stretches of DNA containing modified bases that were inserted during repair. When single-molecule detection of modified bases is used, the sudden transition from all standard bases to mostly modified bases is easily detected using the CCS mode of PacBio sequencing. The method requires that the repair polymerase can readily insert modified bases and that the base modifications do not substantially impair ligation when the missing DNA is replaced and extended to the next stretch of intact DNA. The conditions we have used are effective in this regard, but it is likely other polymerases and conditions could also be used. For detecting the impact of different processing steps, different modified bases could be used at different processing stages.

Because the truncation site can be resolved to a narrow area based on the presence/absence of modified bases, the sequence near the truncation site can be examined for features that might affect its frequency. One obvious feature near the most common truncation sites is the presence of strong hairpins with secondary structure. Short hairpins have been previously observed to lead to AAV truncations (24, 25) so this is not a surprising finding. We have not determined the exact threshold for how much secondary structure is needed for a significant impact because we are most interested in avoiding vector truncations, not enhancing them. Based on the results in Fig 4, the threshold for hairpin stability is approximately -80 kcal/mole over a 150 nt segment. Stronger hairpins will generate frequent truncations while hairpins with less than half that stability will not. We have been able to change vectors to avoid truncations and easily test redesign effectiveness using HIDEN-Seq.

Traditional methodology for testing the quality of AAV preparations for use as gene therapy vectors have clearly shown wide discrepancies in quality. DNA sequencing-based methods have been less widely used, partly because the complexity of the sequence data has complicated its interpretation. Single-molecule, long-read sequencing has provided a better picture of AAV quality (24-28), but data interpretation has still been problematic due to the challenges of working with AAV. Use of the methods described herein and further improvements will make sequence analysis of AAV the standard for quality assessment.

## Methods

All enzymes and non-vector DNAs were purchased from New England Biolabs. DNA sequencing and repair kits were purchased from Pacific Biosciences. Modified nucleotides were purchased from Trilink. Recombinant AAV vectors were produced by triple transfection in HEK293 cells as previously described (28). Genomic DNA was extracted from the vectors by heating at 75C° for 5 min and then purifying DNA via a Monarch PCR and DNA cleanup kit with a 1:2 ratio of sample to binding buffer. DNA was eluted from the column using 11 μl elution buffer. DNA quality was assessed using a HS dsDNA Qubit kit. For many experiments, standard Pacific Biosciences protocols were followed for the End Repair/A-Tailing Step and Adapter Ligation with barcodes. When modified bases were used instead of the standard repair method, the following conditions were used for repair:

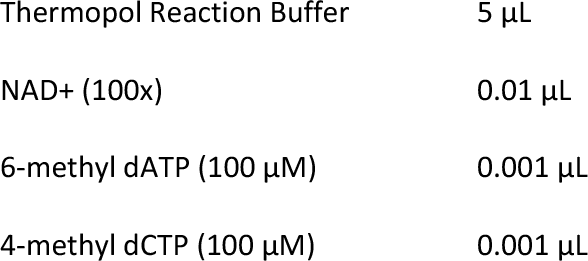
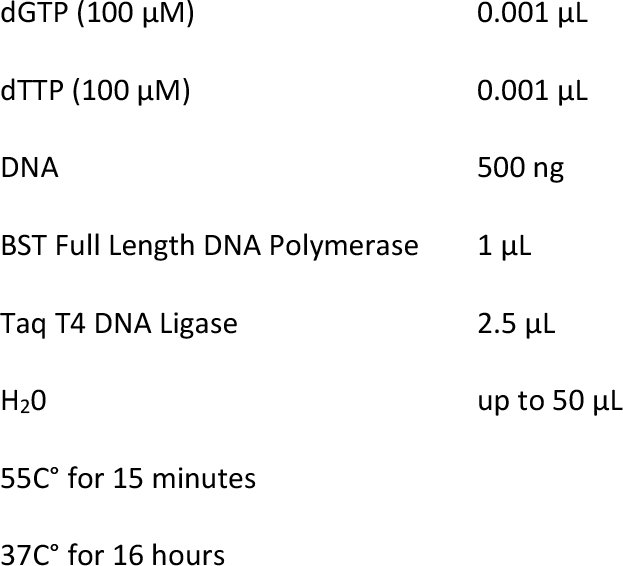

Pronex beads were used to cleanup samples and then pooled for sequencing after barcoding. Standard sequencing was carried out on a Sequel II instrument with a 30 hour movie.

### HIDEN-Seq Analysis

After PacBio SMRT sequencing, HIDEN-seq consists of the following steps: alignment (I), modified base detection (II), and patch identification (III). In the first step, all raw subreads were aligned to a respective reference using the PacBio minimap2 wrapper, pbmm2. Subreads were used rather than CCS reads in order to generate more patch data points for plotting. Aligned subreads were filtered by read length, quality, and for supplementary alignments. In the second step, PacBio’s IpdSummary tool was run on aligned reads to generate files containing IPD ratios for each aligned subread. These IPD summary files were filtered for bases with IPD ratio >3.0, identifying them as modified bases. In the third step, these modified bases were quantified and plotted based on their locations relative to the reference. The plot was then normalized according to CCS read coverage. Looking at the peaks of the plot, we were able to identify regions where modified base repair was occurring. Additionally, through control experiments we were able to determine that 6-methyl-dATP IPD values were more accurate in evidencing modified base repair due to having a higher IPD ratio distribution, compared to 4-methyl-dCTP, which was more susceptible to noise.

